# Genomes contain relics of a triplet code connecting the origins of primordial RNA synthesis to the origins of genetically coded protein synthesis

**DOI:** 10.1101/2021.11.03.467149

**Authors:** Geoffrey H. Siwo

**Affiliations:** Center for Research Computing, Eck Institute for Global Health, Department of Biological Sciences, University of Notre Dame, IN, 46556, USA

## Abstract

Life on earth relies on three types of information polymers-DNA, RNA and proteins. In all organisms and viruses, these molecules are synthesized by the copying of pre-existing templates. A triplet-based code known as the genetic code guides the synthesis of proteins by complex enzymatic machines that decode genetic information in RNA sequences. The origin of the genetic code is one of the most fundamental questions in biology. In this study, computational analysis of about 5,000 species level metagenomes using techniques for the analysis of human language suggests that the genomes of extant organisms contain relics of a distinct triplet code that potentially predates the genetic code. This code defines the relationship between adjacent triplets in DNA/RNA sequences, whereby these triplets predominantly differ by a single base. Furthermore, adjacent triplets encode amino acids that are thought to have emerged around the same period in the earth’s early history. The results suggest that the order of triplets in primordial RNA sequences was associated with the availability of specific amino acids, perhaps due to a coupling of a triplet-based primordial RNA synthesis mechanism to a primitive mechanism of peptide bond formation. Together, this coupling could have given rise to early nucleic acid sequences and a system for encoding amino acid sequences in RNA, i.e. the genetic code. Thus, the central role of triplets in biology potentially extends to the primordial world, contributing to both the origins of genomes and the origins of genetically coded protein synthesis.

**Significance:** One of the most intriguing discoveries in biology is that the order of amino acids in each protein is determined by the order of nucleotides (commonly represented by the letters A, U, G, C) in a biological molecule known as RNA. The genetic code serves as a dictionary that maps each of the 64 triplets ‘words’ in RNA to the 20 amino acids, thereby specifying how information encoded in RNA is decoded into sequences of amino acids (i.e., proteins). The deciphering of the genetic code was one of the greatest discoveries of the 20th century (1968 Nobel Prize in Medicine and Physiology) and is central to modern molecular biology. Yet, how it came to be that the order of triplets in RNA encodes the sequence of the protein synthesized remains one of the most important enigmas of biology. Paradoxically, in all life forms proteins cannot be synthesized without RNA and RNA itself cannot also be synthesized without proteins, presenting a chicken and egg dilemma. By analyzing thousands of microbial genomes using approaches drawn from the field of natural language processing, this study finds that the order of triplets across genomes contains relics of an ancient triplet code, distinct from but closely connected to the genetic code. Unlike the genetic code which specifies the relationship between information in RNA and the sequence of proteins, this ancient code describes the relationship between adjacent triplets in extant genome sequences, whereby such triplets are often different from each other by a single letter. Triplets that are closely related by this ancient code encode amino acids that are thought to have emerged around the same period in the earth’s early history. In other words, a fossil record of the chronological order of appearance of amino acids on early earth appears written in genome sequences. This potentially demonstrates that the process by which RNA sequences were synthesized in the primordial world relied on triplets and was coupled to amino acids available at the time. Hence, the connections between primordial RNA synthesis and a primitive mechanism for linking amino acids to form peptides could have enabled one type of molecule (RNA) to code for the other (protein), facilitating the emergence of the genetic code.

## Introduction

Life’s three most important molecules-DNA, RNA and proteins-are information polymers that encode biological information in their sequences. In all organisms and viruses, the synthesis of nucleic acid sequences predominantly occurs through the copying or replication of pre-existing templates by enzymes. RNA, a molecule widely thought to have preceded DNA, is synthesized by copying of pre-existing DNA sequences except in RNA viruses where it can be synthesized directly from pre-existing RNA templates. The decoding of the genetic information into amino acid sequences is facilitated by the genetic code which maps each of the 64 possible trinucleotides (triplets) to amino acids and stop signals [1–3]. Understanding the origins and evolution of the genetic code is arguably one of the most fundamental and difficult problems in evolutionary biology [3, 4]. This study provides novel observations suggesting that the sequence characteristics of ancient DNA/RNA molecules shaped the structure of the genetic code.

It is envisionable that genomes of extant life may carry signatures of ancient events, the first steps of the evolution of RNA sequences from a limited set of templates that were synthesized from scratch, i.e. *ab initio*, in the RNA world. These ancestral sequences would have been short and random in sequence [5]. Over the past 4 billion years, the initial RNA sequences have been copied and recombined to yield much larger genes and genomes that characterize life. A shared ancestry of natural DNA/RNA sequences from short fragments generated ab initio would imply that diverse genomes are potentially pervaded by sequence signatures that trace back to the early stages of life. However, these sequence signatures would obscured in time due to mutations and recombination events, leaving few clues. Only very short, fundamental and self-reinforcing signatures would be left behind scattered throughout various genomes. The detection of such signatures would not be amenable to conventional phylogenetic techniques used to assess the evolution of nucleic acid sequences. An alternative approach would be to consider this challenge using analogies from the analysis of natural (human) languages.

Genome sequences and natural language texts share many similarities. Both rely on an alphabet from which symbols can be concatenated to encode messages. In any human language, a system of rules or grammar constrains sequences of words that can form grammatical sentences. The evolution of DNA/RNA sequences from the early days of life on earth can be likened to evolution of human languages over time given the symbolic representation and sequential characteristics of both DNA and human language which result in strings of characters drawn from an alphabet. Words in human language consist of letters combined in different ways in a non-random manner. Some combinations of letters (n-grams) are more frequent than others and some words are more likely to occur next to each other than others. In principle, computational techniques primarily developed for the analysis of human language may therefore be useful in extracting complex patterns in biological sequences that may be undetectable by phylogenetic approaches. However, unlike natural language, nucleic acid sequences are not organized into clearly demarcated words. An exception to this is in protein translation where specific consecutive trinucleotides (codons) in mRNA sequences are read by ribosomes with each codon specifying an amino acid. While codons are composed of triplets, they are restricted to triplets that occur in valid reading frames demarcated by the location of a start codon (often AUGs located in specific sequence contexts).

A range of studies open the possibility that a foundational role for trinucleotides in biology predates the emergence of the genetic code. First, trinucleotides and other shorter oligomers can be formed spontaneously under conditions that mimic early earth [6]. Secondly, the use of triplets as building blocks for RNA-catalyzed RNA synthesis by a ribozyme enables it to replicate folded RNA molecules including itself without a primer [7–9], a key requirement for an RNA based origin of life [4, 10, 11]. While ribozymes that can synthesize RNA using mononucleotide substrates have been evolved in laboratory experiments [12–16], their ability to replicate RNA templates including their own sequence is highly impeded by secondary structures [16]. Finally, previous studies have suggested that early tRNA molecules were initially involved in RNA replication using triplets as substrates and were later adopted for protein translation [17–19]. Indeed, the synthesis of proteins in all organisms involves a transient attachment of amino acids to their cognate tRNAs via a triplet sequence (CCA) at the 3-end of tRNA molecules [20, 21]. In this way, even before the emergence of the genetic code, a mechanism for reading RNA triplets probably existed.

To test the hypothesis that trinucleotide patterns across genomes could provide insights into the origins of genomes and the genetic code, this study performed an analysis of triplet patterns in thousands of metagenomes using a technique known as word embedding [22], that captures the relationships between words using numerical vectors. The analysis finds that adjacent triplets of DNA/RNA sequences in diverse microbial genomes have a non-random relationship in their syntax and that this relationship maps to the structure of the genetic code. The detected relationship between adjacent triplets provides insights into potential characteristics of ancient DNA/RNA sequences and the assignment of triplets to amino acids, i.e. the origins and evolution of the genetic code. The results suggest that the origin of the genetic code is intimately connected to an ancient triplet-based synthesis of RNA. This supports the hypothesis that RNA catalyzed triplet-based synthesis of RNA molecules paved the way to a triplet-based genetic code for protein synthesis. In summary, the results imply that the genetic code shares its origin with a triplet-based mechanism for primordial RNA synthesis and is imprinted in the genomes of extant organisms.

## Results

### A language model infers trinucleotide contexts in 4,930 species-level metagenomes

To investigate the potential relationships if any between all 64 trinucleotides in genome sequences, genome-specific language models were generated using one of the largest, high-quality metagenomic datasets consisting of 4,930 species-level metagenome bins [23] (see Materials and Methods for details on dataset). Each metagenome was first split into triplet ‘words’, followed by training of a shallow neural network using a technique known as Word2Vec [22] (Fig. 1). Briefly, given a corpus of text, a language model (here, Word2Vec) learns a vector representation of words in the text such that words that tend to occur adjacent to each other or in similar contexts are represented with similar vectors. The resulting embedding vectors can then be used to predict the context in which words occur, for example to predict the most likely adjacent word given an input word.

**Fig. 1:**
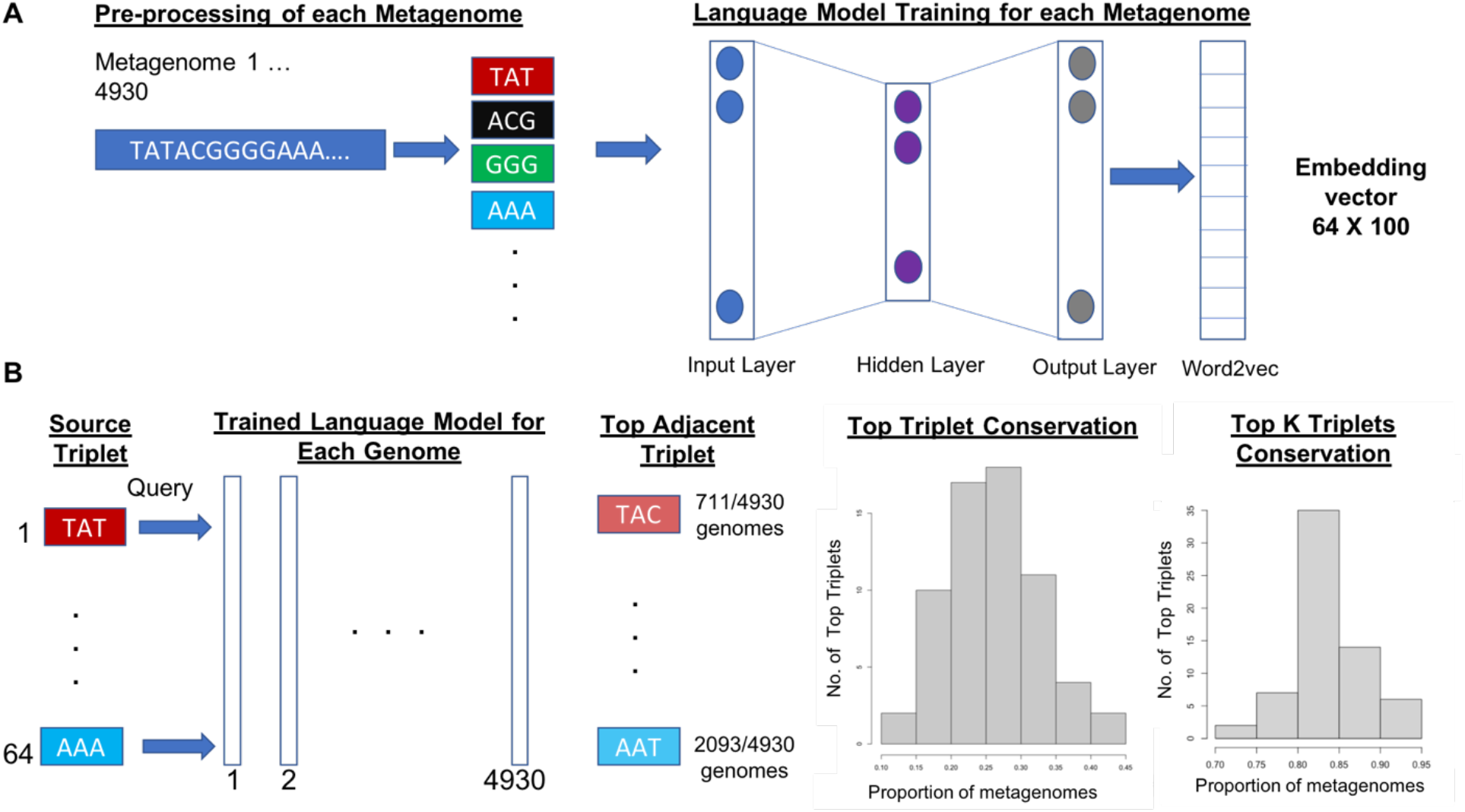
Overview of the language modeling-based approach to inferring triplet contexts. **A**. Training of the language model for each genome using a shallow neural network (Word2Vec) resulting into encoding of triplets in 100-dimensional vector for each of the 64 triplets (64 × 100 embedding space). **B**. Application of the trained language models in inferring triplet contexts for each triplet across all the metagenomes.

For each triplet, a word embedding model was used to infer its context or adjacent triplets in each metagenome. Interestingly, for each of the 64 triplets, the inferred adjacent triplet was conserved across a higher percentage of metagenomes than would be expected by chance (binomial test P < 2.2e-16 for each of the 64 triplets; Supplementary Table 1 and Fig. 1 B histograms). For example, in the case of the triplet TAT whose adjacent neighbor varied the most across genomes, there was a strong preference for a specific triplet (TAC) which was its top inferred neighbor in 14.4% (711 metagenomes) compared to a random likelihood of 0.016 (77 metagenomes), an enrichment by a factor of 9. In contrast, the triplet TTT with the strongest preferred neighbor was enriched by a factor of 27 with the triplet ATT inferred as its top neighbor in 43.2% of cases (2143 out of 4930 metagenomes; P < 2.2e-16; Fig. 1 B). On average, for any given triplet, its top neighboring triplet was conserved in 26.1% of metagenomes (median = 25.4%; binomial test *P* < 2.2e-16), representing an average enrichment factor of 16.4 (median = 15.9). Detailed information for each triplet is provided in Supplementary Table 1.

Furthermore, the observed triplet context of each triplet across metagenomes extend beyond the immediate neighbor. Assuming an equal likelihood of observing any triplet in the top K most frequent neighbors of a given triplet at a P-value threshold of 0.05 (i.e., at least 95 metagenomes), K ranged from 7 to 14 (median = 10; see Supplementary Table 1 for Top K for each triplet). In other words, for a given triplet, each of its top K neighbors are conserved in at least 95 metagenomes. For example, for the triplet AAC, 7 specific triplets (K = 7) were each inferred as its top neighbors at this threshold. Together, the 7 triplets are the top neighbors of AAC in 81% of metagenomes (4034 metagenomes) even though this number of triplets only accounts for 10.9% of triplets. For each triplet, the proportion of metagenomes covered by the top K neighboring triplets ranged from 70.3% to 92.2% (Fig. 1 B; Supplementary Table 1).

### Trinucleotides occurring in similar contexts are related in sequence

To visualize the connections among all the triplets based on their top adjacent neighbors, a directed network was constructed in which each node represents a specific triplet and edges connect one triplet (‘source triplet’) to another triplet (‘target triplet’) if the target triplet is inferred as the top neighbor of the source triplet. A visualization of the resulting network is shown in Fig. 2.

**Fig. 2.**
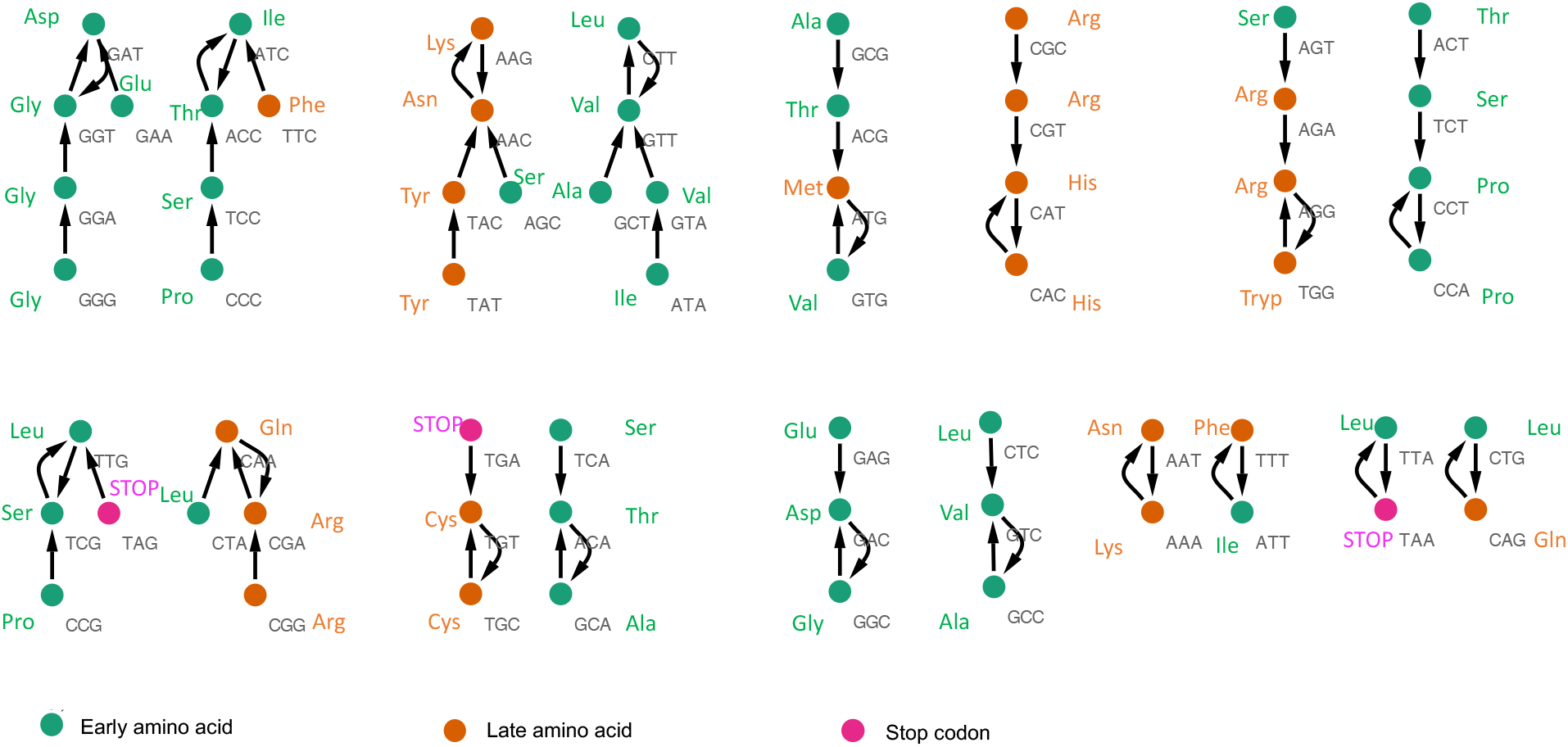
Visualization of the triplet network showing that each triplet is connected to a triplet from which it differs by a single base. The nodes are colored based on whether the triplet corresponds to a codon for an early or late amino acid or stop codon. Classification of amino acids as early or late is based on the inferred abundances on early earth using a consensus of observations from on prebiotic chemistry experiments, isolation of amino acids from meteorites, phylogenetic studies and thermodynamic simulations.

Remarkably, examination of the network shows that each triplet is connected directly to a triplet that differs from the source triplet by a single base (Fig. 2). For example, the triplet *GG****G*** is directly connected to *GG****A*** which is connected to *GG****T***; *G****G****T* is connected to *G****A****T* and *GA****T*** is connected to *GA****A*** (with the nucleotide that differs between consecutive triplets highlighted in bold).

In order to assess the likelihood of a network composed of 64 triplets in which all connected triplets are 1-base change away from each other, we performed 10,000 simulations of randomly connected sets of 64 triplets, whereby each triplet is randomly paired with another triplet (Fig. 2). In none of the random sets was a set of 64 triplets obtained in which each triplet is paired with another triplet 1-base change away, demonstrating that the likelihood of randomly observing such networks is low (P < 0.0001). Indeed, within the 10,000 randomly generated triplet sets, the maximum number of triplets in a set of 64 that are 1-base change away from each other was only 24 and occurred only once (P = 0.0001). In contrast, the minimum number of triplets in 10,000 random sets of 64 that are 1-base change away was 1 and the median was 9 triplets. Randomly connected triplet pairs in the simulated networks were more likely to be 2 or 3 mutation steps from each other with a median of 27 out of 64 triplets being 2 or 3 base change away in each of the 10,000 random networks.

The triplet network above is constructed only by considering for each triplet, its top inferred neighbor. To understand whether the observed syntactic relationships between triplets in the network extends beyond the top neighbors, for each triplet an assessment of its sequence similarity to its top K neighbors observed at a threshold of 95 genomes (P < 0.05) was performed. At this threshold, the number of consecutive triplets that are 1-base change away from a source triplet ranged from 2 to 9 with a median of 5 triplets. In other words, most top K triplets occurring in greater than 95 metagenomes (P= 0.05) for each triplet are 1 base change away from the source triplet implying that neighboring triplets were likely derived from the same parental triplet. For example, assuming the ancestral sequences were rich in triplet repeats, accumulation of mutations could lead to the observed pattern.

These results show that triplets have a preferred context i.e., the likelihood of observing a given triplet next to another triplet is not equally likely and extends beyond immediate neighbors. This is especially notable given that the language model used to infer the triplet relationships was trained indiscriminately on metagenome sequences that include non-coding sequences. Furthermore, model training was not restricted to open reading frames (ORFs). Thus, it is conceivable that the observation for a strong preference between neighboring triplets is not due to codon bias, a well-studied phenomenon that reflects the preferences for certain synonymous codons within ORFs [24–26]. Given the very low likelihood of observing the above pattern of triplets and its detection across diverse metagenomes, it is likely that it reflects an ancient pattern of common ancestral sequences of diverse microbial species. Henceforth, this study refers to this pattern as a triplet code.

### Triplet code network recapitulates symmetries in genomic DNA single-strands (Chargaff’s second parity rule)

The triplet code network shown in Fig. 2 self-organizes into 18 distinct tree-like structures or connected components. A close examination of the structures reveals that each has a similar topology to at least one other structure. The 18 structures can thus be organized into 9-pairs in which members of a pair have the same topology (Fig. 2). Remarkably, each pair of clusters that have a similar topology consists of triplets that are the reverse complements of each other arranged in a similar manner. For example, the structure containing the sequence *TGA-TGT-TGC* has the same topology as its reverse complement *GCA-ACA-TCA*.

This observation is surprising because the language model used to generate the triplet code network was only trained on single-stranded genomic DNA sequences yet it appears to have learned the base-pairing complementarity of nucleotides and the polarity (5′→3′ directionality) of nucleic acid sequences.

The complementarity between nucleotides underlies base pairing which would seem obvious when considering double stranded sequences [27]. However, before the discovery of the structure of DNA by Watson and Crick, one of the main clues leading to this discovery was the observation by Erwin Chargaff that in natural DNA sequences, A and T were always at a 1:1 ratio while G and C were also in a 1:1 ratio [28–30]. This observation commonly known as Chargaff’s first parity rule would seem trivial in hindsight because of knowledge of base-pairing. Unexpectedly, in the analysis of single strands of DNA in almost all extant genomes, it has also been found that the frequency of any nucleotide or short oligonucleotide (∼10 nts) is approximately equal to the frequency of its reverse complement, a phenomenon referred to as Chargaff’s second-parity rule [30–34].

While Chargaff’s first rule is resolved, the second rule remains mysterious in origins and function, and cannot be accounted for by base-pairing. The triplet code network reflects that the language model captures Chargaff’s second parity rule. This is perhaps the first demonstration that the second parity rule holds when considering the contextual relationships between trinucleotides, going beyond a frequency-based rule.

### Triplet code is connected to the structure of the genetic code and the chronological abundance of amino acids on early earth

The proposed triplet code defines the relationship between neighboring triplets which may reflect a combination of mutational signatures or physicochemical constraints in nucleic acid sequences. It is also possible that in addition to these factors, the triplet code may represent relics of ancient DNA/RNA de novo synthesis or template-dependent (replication) mechanisms. Since the decoding of genetic information during RNA translation also proceeds using codons that map the sequence of triplets to amino acids, it is conceivable that constraints on the order of triplets in RNA sequences in the primordial world would also constrain the assignment of amino acids to specific triplets (codons), i.e., the emergence of the genetic code. This is especially so because the availability and abundance of specific amino acids has changed dramatically during the earth’s early history with simple amino acids such as glycine envisioned to have emerged early in the prebiotic era and more complex ones such as tyrosine later [35–38].

Thus, to investigate potential relationships between the topological connections of the triplets, the structure of the genetic code and chronology of amino acids, each node in the triplet network was also labelled by the encoded amino acid based on the standard genetic code (Fig. 2). Each node was also color coded by a consensus [35, 37, 38] of whether the assigned amino acid has been consistently produced in prebiotic chemistry experiments [39–42], and has been detected in meteorites (‘early amino acid’) [43, 44] or emerged later (‘late amino acids’). Furthermore, studies show that the concentration of the early amino acids in the recent evolution of proteins is mostly decreasing while that of late amino acids is mostly increasing [35, 36]. In addition, the thermodynamically cheapest amino acids are among the early ones [45].

Integration of this classification scheme into the triplet network shows that triplets corresponding to early amino acids are more likely to be connected to those of other early amino acids than to late amino acids and vice versa (Fig. 2). Out of 22 triplets that correspond to late amino acids, only 32% (7) are connected to triplets of early amino acids with 68% (15) connected to triplets of other late amino acids (binomial test P = 0.0001). In contrast, out of 39 triplets corresponding to early amino acids, 82% (32) are connected to triplets of other early amino acids while 18% (7) are connected to those of late amino acids (binomial test P = 0.02). Two triplets for stop codons are connected to those of early amino acids while the remaining stop codon triplet is connected to a triplet of a late amino acid (Fig 2). Among the 7 triplets for a late amino acid connected to an early amino acid, 2 correspond to phenylalanine codons (TTC and TTT), another 2 to glutamine codons (CAA and CAG), 1 for arginine (AGA), 1 for methionine (ATG) and 1 for asparagine (AAC). The triplets for the early amino acids Gly, Ala, Asp, Gly, Glu and Pro are only connected to other early amino acids. Early amino acids with at least 1 of their triplets connected to that of a late amino acid are: Thr (ACG), Ile (ATC, ATT), Leu (CTA, CTG), Ser (AGT, AGC) and Val (GTG).

It is possible that at some stage in the early evolution of the genetic code, triplets corresponding to codons for specific early amino acids were reassigned to or captured over time by late amino acids. This reassignment may therefore be reflected by triplets of early amino acids connected to those of late amino acids. Consistent with this, the start codon ATG for methionine (Met), considered to have been a key codon capture event from the Ile (an early amino acid) is connected to a triplet for another early amino acid (Val). However, in the previous studies the methionine codon capture was proposed to involve capture of Ile triplets [46–48] while the results in this study open the possibility of capture from GTG (Val) or ACG (Thr). Interestingly, GTG and ACG are one of a few codons that can be read occasionally as start codons in the place of ATG (Met) [49], with GTG being used in as much as 14% of genes in *E. coli* [50]. Notably, from the triplet network in this study, even though Ile triplets ATC and ATT are not directly connected to ATG, they are among the few early amino triplets that are connected to those of late amino acids. Furthermore, in agreement with a previous study suggesting that the late amino acids Phe, Gln, Met and Asn captured codons from other amino acids before LUCA [51], triplets for these amino acids are connected to those of early amino acids.

### Triplet code reconstructs a key translation initiation signal in prokaryotes

It is possible that the non-random association between a given triplet and its top K neighbors may capture sequence signatures that impact decoding of the triplet. Out of the 64 possible triplets, the start codon (ATG) is unique in that it is the only codon that, even though in all organisms specifies a single amino acid (methionine), is decoded by a different tRNA depending on its sequence context. When located at specific sequence contexts demarcating the start of an ORF, ATG is decoded into methionine by an initiator tRNA (fMet-tRNA in prokaryotes) compared to its decoding by a different tRNA (tRNA-methionine) when located at internal sites. In bacteria and archaea, a critical signal for translation initiation consists of a short sequence known as the Shine-Dalgarno sequence (SD, ribosome binding site), typically located 11 to 13 nucleotides from the start codon [52]. 16s rRNAs of many prokaryotes contain a sequence complementary to the SD sequence. Several studies have shown that this anti-SD sequence facilitates the recruitment of 30s ribosomal subunit onto mRNA and the subsequent alignment of the start codon with the ribosomal P-site [53–58].

To explore whether the top K neighbors of the triplet ATG reflect sequence signatures associated with translation, the top K triplets of ATG were concatenated into a continuous string (Fig. 3). Assuming that ATG is at the 3’-end of the resulting sequence, the sequence motif matching the SD consensus (GGAGGT) occurs 11 nucleotides upstream of the ATG (Fig. 3). This observation is intriguing especially given that the triplets starting from the −1 position of the sequence and ending at the −10 position are all 1-base change away from ATG. Furthermore, triplets nearest to ATG (i.e., GTG and ACG) are common alternative start codons [49, 59]. Thus, this suggests that syntactic relationship observed between adjacent triplets could have facilitated the emergence of the SD sequence in the context of ATG.

**Fig. 3.**
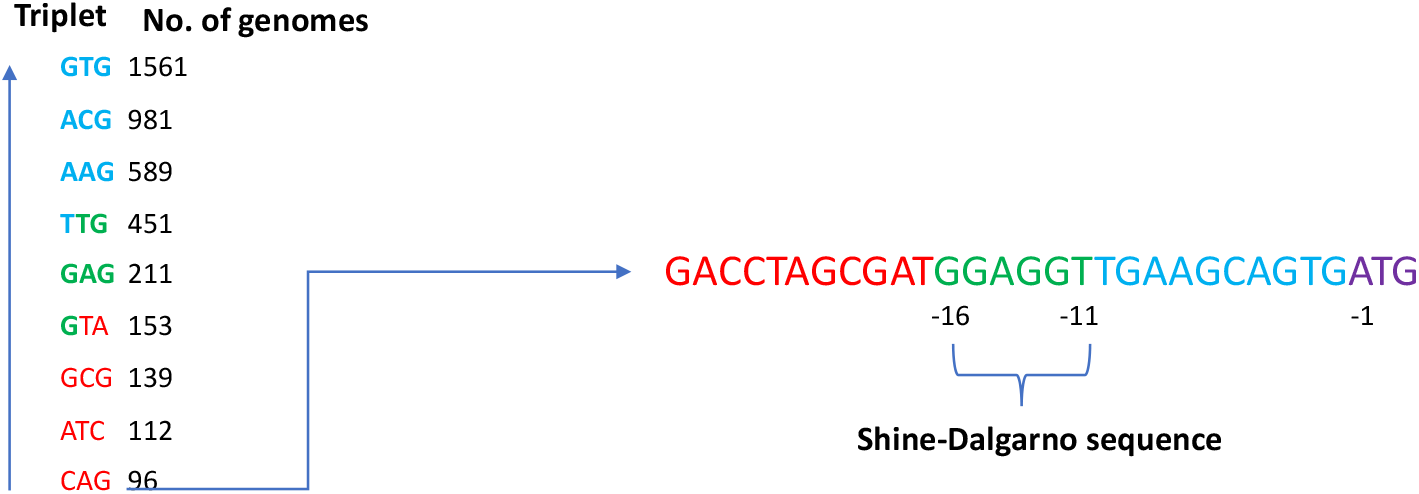
Analysis of the context of the ATG triplet shows a Shine-Dalgarno (SD) consensus sequence is located 11 nucleotides away, approximating the observed location of the motif in bacterial genomes.

The reconstruction of an SD-like sequence using the triplet patterns learned by a language model has important implications in understanding the origins of translation because anti-SD sequences are present in 16s rRNAs of many bacterial and archaeal species. Furthermore, while the language models in this work were trained without being provided with explicit information on ORFs and non-coding regions, the reconstruction of an SD-like sequence which is located in 5’-untranslated regions (UTRs) of mRNAs shows that signals located in non-coding regions can co-evolve with those that are decoded into amino acids. It is notable that while not all ATG triplets are located in the context of SD motifs, and in fact translation in prokaryotes can be initiated in some mRNA sequences that lack SD sequences [60–62], the ability of the language model to infer the association between ATG and SD sequences implies that the two are tightly linked. As anti-SD sequences occur in 16s rRNAs that were likely present in the pre-LUCA period, it is possible that the coevolution of ATG and SD sequences dates to billions of years ago.

## Discussion

In all organisms, the genetic code is the major bridge between genetic information (DNA/RNA) and protein synthesis. While the genetic code plays a critical role in the synthesis of proteins, why and how genetically coded protein synthesis emerged early in the evolution of life is one of the most challenging questions of modern biology [3, 4, 63]. Even though the ability of RNA to serve as a purveyor of genetic information, replicase and catalyst have made it the most widely considered as present at the origins of life [16, 64–66], the synthesis or replication of long RNA molecules has not be realized by RNA alone and requires proteins which are genetically coded. This chicken and egg dilemma leaves room for the possibility that the intimate relationship between the synthesis of RNA and protein sequences emerged in the primordial world, potentially preceding the emergence of the genetic code or as suggested by this study, facilitating its emergence.

Given that the genetic code represents an assignment of each triplet to a specific amino acid, it is conceivable that its emergence was constrained by the order of triplets in primordial RNA sequences and the availability of specific amino acids. The order of triplets in primordial sequences initially synthesized *ab initio* and replicated by non-enzymatic processes was likely semi-random, assuming different chemical reactivities and concentrations of individual nucleotides alongside variable stability and replication fidelity of different sequences. Furthermore, it is possible that triplets were a fundamental unit of primordial RNA synthesis and replication [7, 8, 17, 18, 67].

Based on the above arguments, it can be hypothesized that natural genome sequences are imprinted with triplet-based signatures that reflect the origins and evolution of the genetic code. In 4 billion years of evolution, most sequence signatures from the primordial world have been lost in time by the processes of mutation and natural selection. However, because most naturally existing DNA/ RNA sequences are not synthesized *de novo* but through replication, ligation and recombination of pre-existing templates, it is possible that short sequence signatures that pervaded primordial sequences remain preserved, albeit scattered within and across genomes of diverse species. Furthermore, such signatures may be fixed in genomes and self-perpetuating if they are tightly linked with the integrity of the genetic code or intrinsic error profile of replication or the physicochemical constraints in DNA/RNA.

### Implications of the triplet networks on primordial sequences and the earliest genomes

The results presented in this study showing that each of the 64 triplets have strong preferred sequence contexts characterized by close syntactical relationships of adjacent triplets that encode amino acids with shared evolutionary histories suggests that the origins and evolution of the code is imprinted in natural genome sequences. These observations imply that primordial sequences were likely rich in triplet repeats that overtime accumulated point mutations leading to the observed pattern. Indeed, previous studies have reached a similar conclusion through distinct arguments [68–70]. Assuming mutations are randomly distributed, they are not sufficient to erase the triplet pattern because adjacent triplets have an equal chance of undergoing mutations. In addition, there are only a limited number of mutations that can occur in each triplet, including back mutations. For mutations to totally erase the triplet code, they would need to occur disproportionately in all adjacent triplets.

It is notable that the topology of the connected components of the triplet network is consistent with Chargaff’s second parity rule even though the network was generated using only the positive strand of sequenced metagenomes used in this study. This suggests that base-pairing and the polarity of double-stranded DNA sequences is implicitly stored in single-strands of DNA, supporting the idea that genomes evolve by transpositions and inversions [71–75].

Thus, a simple process through which the earliest genomes (‘pre-genomes’) could have arisen could be through mutations occurring in short RNA sequences rich in repeats that underwent single point mutations to generate a diverse set of sequences. These sequences could have recombined and ligated to generate much longer sequences.

### A shared, non-Darwinian origin of primordial RNA synthesis and genetically coded protein synthesis

The observed association between the context of triplets in genome sequences and the structure of the genetic code is not limited to protein coding regions of genomes. Hence, the results presented are of significance in understanding the origins of genomes and the origins of the genetic code. In principle, the conserved context of triplets in genomes can either be due to constraints arising from the physicochemical properties of different triplets and/or constraints associated with the primordial RNA synthesis. However, even if physicochemical properties of triplets would have an effect on the observed patterns of triplets they cannot account for the connection between triplet contexts and the chronological appearance of amino acids. One possible mechanism through which the structure of the genetic code became imprinted across natural genome sequences independent of open reading frames could be due to primordial synthesis of RNA involving linkage of trinucleotides coupled to amino acids. In this arrangement, two trinucleotides each linked to an amino acid would facilitate an energetically favorable reaction in which the formation of a peptide bond between the adjacent amino acids brought together on a template provides the energy required to ligate the released trinucleotides to each other, resulting in the co-synthesis of RNA and peptides (Fig. 4). Such a process could remove the need for the independent emergence of RNA and protein synthesis, and could have facilitated rapid transition to a ribonucleoprotein world in which the two molecules are synthesized in parallel. Furthermore, if triplets were substrates for both peptide and RNA synthesis in the primordial world, one can argue that this would favor the later emergence of a triplet based genetic code with a ribosome that ratchets 3-nucleotides at a time. This process would be initially non-Darwinian in the sense that primordial sequences would not need to explore alternative genetic codes (e.g., based on dinucleotides instead of trinucleotides or random codon assignments) or need selective pressure to emerge. Thus, the code’s initial origin would be non-Darwinian but its refinement and increasing complexity including use of tRNAs would occur by evolutionary processes.

**Fig. 4.**
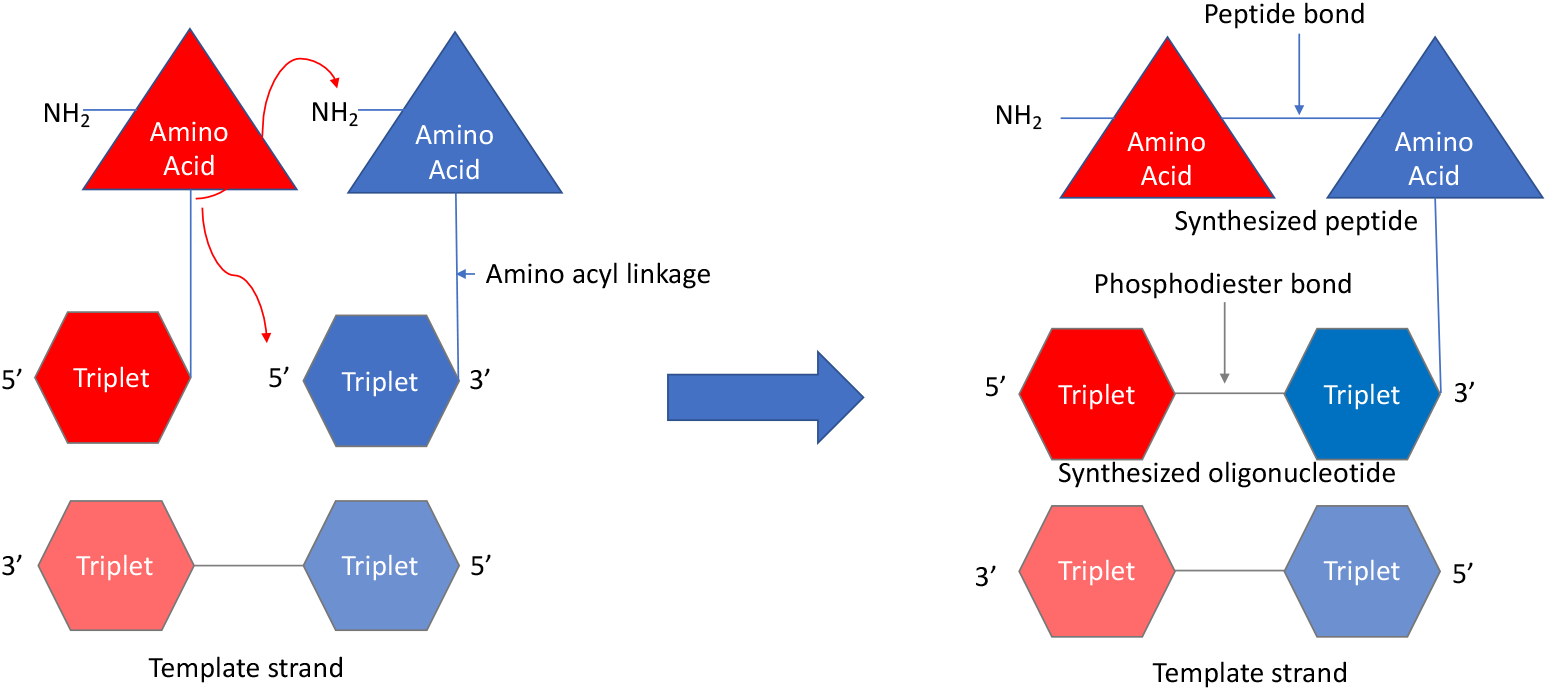
Proposed coupled, templated RNA and peptide synthesis.

The ideas presented above share some characteristics with a number of hypotheses previously put forward on the origins of genetically coded protein synthesis, in particular by Sutherland and Blackburn who proposed that it could have evolved from coupled RNA and peptide synthesis involving amino acids linked to triplets [67]. One of the major challenges to this hypothesis is that in the absence of specific enzymes or catalysts, it is not clear how amino acids could become charged (linked covalently) to specific triplets. One possibility is that stereochemical affinities and chemical reactivity between specific triplets and amino acids could favor the coupling of the specific triplets to amino acids. This would be consistent in part with the stereochemical theory of the genetic code origins proposed by Woese and others [51, 76–83] with a key difference being the additional need for chemical reactivity between amino acids and triplets. In theory, triplets that with a high reactivity to a given amino acid could ‘capture’ the amino acid irrespective of their stereochemical affinities for each other. If such triplets also formed highly labile aminoacyl ester linkages to the specific amino acid, spontaneous hydrolysis of the linkage could provide energy for the coupled formation of more stable peptide and phosphodiester linkages (Fig. 4). The co-synthesis of RNA and peptides could then result in RNA sequences linked to their encoded peptides, enhancing their co-evolution through a genotype-phenotype linkage. Indeed, covalent linkages between mRNAs and the proteins they code for has been used as a strategy for directed evolution in techniques such as mRNA display [84].

Protein synthesis in extant organisms involves the formation of a covalent linkage between amino acids and a trinucleotide sequence (CCA) at the 3-end of tRNAs [20, 21]. While this reaction is facilitated by aminoacyl tRNA synthetases which would have been absent in the primordial world, the fact that in all organisms, peptide bond formation occurs when the attached amino acid is released from the CCA end emerged pre-LUCA. Out of all the trinucleotides, simulations of aminoacyl ester linkages to different triplets shows that CCA aminoacylation results in the highest solvent exposure of the aminoacyl ester bond and a weaker dependence of such exposure on amino acid side chain interactions [85]. It is therefore possible that a simpler system in which peptide synthesis occurs using directly aminoacylated, free trinucleotides (without tRNAs) were replaced over evolutionary time with universal ancestral tRNAs terminated by CCA. It is interesting to note that CCA codes for proline which, compared to other amino acids, forms the least stable aminoacyl ester linkage with its tRNA [86].

### A testable hypothesis for primordial RNA synthesis coupled to ribosome-free translation

The proposed coupled synthesis of RNA and peptides requires reactions that can proceed without the involvement of RNA replicating enzymes/ ribozymes or protein synthesis machinery including rRNAs, tRNAs and ribosomes. Experimental validation of the hypothesis will require testing the hydrolytic stability of amino-acyl linkages between all the triplets and their cognate amino acids and whether such linkages can drive RNA and peptide synthesis on a template. The feasibility of such experiments is supported by at least two studies. First, templated, non-enzymatic RNA synthesis can occur through several mechanisms including through ribozymes [7, 9, 12–16] or using activated mono- or short oligonucleotide substrates [87–90]. The latter is more relevant to RNA synthesis in the primordial world before the emergence of RNA replicases. In relation to this, the Szostak lab recently demonstrated that the presence of a 3-amino group at the 3’-end of tetranucleotides enhances their ligation in the presence of an N-imidazole organocatalyst [90]. Secondly, it has been recently shown that coded peptide synthesis can proceed without tRNAs or ribosomes, specifically using single nucleotides charged with amino acids [91, 92]. One can expect that charged trinucleotides may form more stable base-pairing interactions with corresponding templates compared to those observed with mononucleotides. The proposed coupled synthesis can be viewed as combining these two independent studies into one in which aminoacylated trinucleotides are used to simultaneously drive templated synthesis of RNA and peptides.

It is important to highlight several limitations of this study. First, events that occurred billions of years ago are in a sense unknowable. Furthermore, the observed triplet patterns may have arisen due to other factors that are not considered in this study as there are many unknown factors on the origins and evolution of life. Finally, the analysis presented in this work is based on microbial sequences from human metagenomic studies which may miss important insights from unculturable environmental microbes and extinct microorganisms. However, preliminary of environmental metagenomes shows that the observed triplet code is conserved (data not shown).

Based on the presented finding, this study concludes that the triplet code is a relic of ancient sequences that date back to the pre-LUCA period, potentially to the primordial world. The results open the possibility that coupled primordial RNA and peptide synthesis involving aminoacylated triplets favored the emergence of genetically coded protein synthesis mediated by a triplet-based code. In summary, the work has implications for understanding the nature of ancient DNA/RNA sequences, the origin and evolution of the genetic code and the potentially constrained nucleic acid sequence space explored in nature.

## Materials and Methods

### Metagenome Datasets

Microbes have enormous diversity relative to eukaryotes and are critical in terms of understanding the early evolution of life on earth. Hundreds of thousands to millions of microbial genomes have been sequenced to date and in principle the analyses presented in this study could be applied to any of the available genome sequences. However, publicly available genome sequences have high variability in quality and also contain many redundancies, for example, multiple sequences of the same microbial species. Furthermore, most microbes remain unculturable resulting in skewed representation of sequenced and assembled genomes towards culturable microbes. Metagenomics sequencing can lead to the identification of diverse microbial species including those that are unculturable. Unfortunately, most metagenomics studies are noisy and consist of unassembled genomes. Informed by these issues, this study selected one of the largest, high-quality metagenome datasets that includes 154,723 microbial genome assemblies and 4,930 species-level metagenome bins obtained from 9,428 human metagenomes [23]. The metagenomes spanned multiple body sites, ages, lifestyle and geography. The assembled genomes included 3,796 species level clades drawn from 34, 205 genomes) that had not been sequenced previously. Both bacterial and archaeal genomes are represented in the dataset.

### Training of the language (embedding) models

To analyze trinucleotide (triplet) patterns across the 4,930 species-level metagenome bins, word embedding [22]-a technique commonly used in the analysis of natural language was used. Briefly, word embedding is a computational approach for representing words using numerical vectors that capture the relationships between words based on their co-occurrence or context. A popular technique used for word embedding is Word2Vec, first proposed by Mikolov et al [22]. Words that occur in the same contexts or are adjacent to each other within a corpus of text have similar embeddings.

Typically, natural language text is used as input to word2vec. Unlike natural language, nucleic acid sequences do not have words per se. In this study, we considered triplets as words given their role in protein coding genes as codons as well as their possible roles in primordial RNA synthesis as discussed in the introduction. In addition, the use of triplets as words during language-based modeling of genome sequences by previous studies has demonstrated that they capture important biological information that can be used for different prediction tasks such as the inference of DNA replication origins [93], identification of enhancers [94] or transcription factor binding sites [95]. Therefore, to apply word2vec to the metagenomes, each species level metagenome sequence was split into non-overlapping triplets, starting from the 5’-end. Only one strand (the positive strand) was used in all analyses and no considerations of open reading frames (ORFs) or coding vs. non--coding sequences were made. Each of the metagenomes processed in this way was used as an input corpus to the Word2Vec algorithm implemented in the *gensim* package of the Python programming language.

As an example of how word vectors were obtained for each genome, for example given a genome sequence “TATACGGGGAAA…” the following steps were undertaken (Fig. 1):

- First each genome was split into non-overlapping triplets “TAT ACG GGG AAA …”. In previous biological sequence analysis tasks [93, 94], overlapping K-mers were used during word embedding. However, in this study the use of overlapping K-mers would introduce a statistical artefact in examining the relationships between adjacent triplets since overlapping triplets obtained by a sliding through the sequence 1-nucleotide a time would be predictably different by 1 nucleotide. Hence, pre-processing the sequences in such a way would introduce strong correlations between adjacent triplets.
- To learn a vector representation of each triplet, Word2Vec was then trained on the split sequence using an approach known as Continuous Bag of Words (CBOW). CBOW learns a vector representation of each word (triplet) by considering the adjacent context words in a specified window size to predict the current word. An alternative approach is known as the Skip Gram in which the model is trained to predict the context given the adjacent words. CBOW was used in this study for its fast speed.
- A window size of 1 was used to define the number of adjacent words to consider. At this window size given the split sequence above, to predict the word ACG using CBOW, 1 word to the left (TAT) and 1 word to the right of ACG (GGG) is considered. Because given any genome sequence, each of the 64 triplets occurs multiple times, the embedding vector for each triplet was obtained by learning the weights that maximize the probability of predicting the triplet through backpropagation in a shallow neural network shown in Fig. 1
- The number of dimensions used to represent each triplet was set to 100 and a minimum count for each triplet to be considered in the model was set to 10.
- Embedding models were trained for each genome separately resulting in 4,930 separate models that were stored for further analysis in a KeyedVector format which provides a mapping between each triplet and its corresponding vector.

The language model files are provided as part of the online Supplementary Files on Github.

### Predicting adjacent triplets using the trained language models for each genome

To predict the top adjacent triplet for each triplet in each genome, the KeyedVectors for each metagenome was queried in *gensim*. For each metagenome, the top adjacent triplet for each triplet was defined as the triplet whose embedding vector has the highest positive similarity to the source (query) vector based on cosine similarity. Given each triplet, the frequency at which a single specific triplet was its top neighbor across the metagenomes was then computed.

To construct a directed network showing relationships between triplets, each query triplet (node) was connected to a triplet that is its top neighbor across the metagenomes. The edges in the network represent the frequency at which a given source triplet is adjacent to the connected triplet. The network visualization was performed using Cytoscape [96].

### Statistical assessments

For each triplet, the statistical significance for observing a specific triplet as its top neighbor in *n* out of 4930 metagenomes was computed using a binomial test in R (*binom*.*test(n,4930,0*.*015625)*), assuming a probability of 1/64 (0.015625) of each of the triplets to have equal likelihood to be the most frequently adjacent to a given triplet. This also assumes independence between species-level metagenomes based on species definition.

To determine the top K triplets for each triplet at P = 0.05, and assuming equal likelihood of occurrence of any triplet as the top K^th^ neighbor, a binomial test was used. Under these assumptions, for each triplet, the top K^th^ triplet was determined by sorting the inferred neighbors by the number of metagenomes in which they are inferred as the top neighbor of the source triplet with a threshold of 95 metagenomes.

The statistical significance of observing a top triplet that is one base change away from the target (source) triplet was estimated by determining the number of triplet pairs that are one-base changes away from each other in random 10,000 “genetic codes” each containing 64 triplets.

### Selection of early and late amino acids

The chronological order of appearance of amino acids on early earth was based on a consensus [35] of previous studies including the Miller-Urey experiments [39–42] and studies on amino acids isolated on meteorites [43, 97]. The 10 early amino acids were based on prebiotic chemistry experiments and also have been identified in meteorites in the following order of relative abundance: Gly, Ala, Asp, Glu, Val, Ser, Ile, Leu, Pro, Thr [35]. This is also in agreement with consensus approaches for the recruitment of amino acids into the genetic code [37, 38] and correlates with expectation of free-energy synthesis of different amino acids [45].

## Acknowledgements

I am grateful to Dr. Arvind Varsani (Arizona State University) and Dr. Michael Ferdig (University of Notre Dame) for comments on this manuscript.

## Supplementary Materials

Supplementary Table 1 and additional information are available on GitHub at https://github.com/SiwoResearch/Triplet_Code

